# An organotrophic *Sideroxydans* reveals potential iron oxidation marker genes

**DOI:** 10.1101/2025.02.27.639646

**Authors:** Rene L. Hoover, Kirsten Küsel, Clara S. Chan

## Abstract

To understand the ecophysiology and the role of iron-oxidizing bacteria (FeOB) in various ecosystems, we need to identify marker genes of the iron oxidation pathway to track activity *in situ*. The Gallionellaceae *Sideroxydans* sp. CL21, an autotrophic iron-oxidizing bacteria isolated from a peatland, is unusual amongst FeOB isolates in its genomic potential to utilize organic compounds as energy sources. Therefore, it offers the unique opportunity to determine genes expressed under litho- versus organotrophic conditions. We demonstrated the growth of *Sideroxydans* sp. CL21 on organic substrates (lactate and pyruvate) and inorganic substrates (Fe(II), magnetite, thiosulfate, and S(0)). Thus, cells were capable of lithoautotrophic, organotrophic, and potentially organoheterotrophic growth. Surprisingly, when lactate-grown cells were given Fe(II), mid-log phase cells were unable to oxidize iron, while late-log phase cells oxidized iron. To elucidate iron oxidation pathways, we compared gene expression between mid-log (non-iron-oxidizing) and late-log (iron-oxidizing) lactate-grown cells. Genes for iron oxidases (*cyc2, mtoA*) were highly expressed at both time points, so did not correspond to iron oxidation capability, making them unsuitable marker genes of iron oxidation activity by themselves. However, genes encoding periplasmic and inner membrane cytochromes were significantly upregulated in cells capable of iron oxidation. These genes include *mtoD*, *cymA/imoA*, and a cluster of Fe(II)-responsive genes (*ircABCD*). These findings suggest Gallionellaceae regulate their iron oxidation pathways in multiple stages, with iron oxidase-encoding genes proactively expressed. Other genes encoding electron carriers are upregulated only when iron oxidation is needed, which makes these genes (i.e. *ircABCD*) good prospective indicators of iron oxidation ability.

**Importance:** FeOB are widespread in the environment and we suspect that they play key roles in nutrient and other elemental cycles. However, with no isotopic marker, we lack the ability to monitor FeOB activity, prompting us to search for genetic markers. Previous work suggests that expression of iron oxidase genes does not directly correspond to iron oxidation activity in Gallionellaceae and little was known about the other genes in the pathway. Here we study a unique FeOB isolate that possesses organotrophic capabilities and demonstrate its potential for mixotrophic growth on lactate and Fe(II). Its ability to oxidize iron is regulated, allowing us to discover potential iron oxidation pathway genes with expression that corresponds to iron oxidation activity. If these genes can be further validated as iron oxidation marker genes, they will enable us to delineate autotrophic and organoheterotrophic FeOB impacts on carbon cycling in wetlands and other natural and engineered environments.

## Introduction

Iron oxidation is a widespread process in the environment, playing key roles in water and soil health as Fe(III) oxyhydroxides can adsorb various nutrients and metals (1). Despite the pervasiveness and importance of iron oxidation, there remains much uncertainty as to how much of it is abiotic versus biotic, i.e. catalyzed by iron-oxidizing bacteria (FeOB). This distinction (biotic vs. abiotic) is key to understanding how iron oxidation affect other nutrient cycles, which requires 1) a deeper understanding of C metabolisms of FeOB and 2) a method to link microbial iron oxidation activity in the environment to C and other metabolic transformations. Although FeOB are widely thought to be primarily autotrophic, accumulating evidence suggests a considerable number of FeOB may be mixotrophic (2–6) or organo(hetero)trophic (7, 8), expanding their potential effects on C cycling. Because these FeOB can use alternate energy sources (9–12), they are not always oxidizing iron, so we require a method of detecting active iron oxidation. To link microbial iron oxidation to biogeochemical effects, we need gene markers of iron oxidation activity.

Studying facultative FeOB allows us to discern genes specific to iron oxidation. An isolate of the Gallionellaceae, *Sideroxydans* sp. CL21, presents a new opportunity to learn about FeOB metabolism, iron oxidation mechanisms, and potential marker genes. *Sideroxydans* sp. CL21 is rare amongst FeOB isolates in possessing genes for organoheterotrophy, notably lactate permease and dehydrogenase. It was isolated from the Schlöppnerbrunnen fen (13), a semi-acidic (pH 4.5 to 5.5), iron-rich peatland in Northern Germany (14–16). *Sideroxydans* sp. CL21 is closely related to the well-studied *Sideroxydans lithotrophicus* ES-1 (9), but differs from ES-1 in that its genome is ~20% larger and encodes genes for organoheterotrophy (17, 18). *Sideroxydans* sp. CL21 has been shown to grow on the non-iron substrates H_2_ and thiosulfate (18). It has also been grown on the reduced iron substrates FeS and Fe(0) (13, 18), but neither of these are pure Fe(II) substrates as Fe(0) evolves H_2_, so growth on Fe(II) alone has not been demonstrated. *Sideroxydans* sp. CL21 has also been grown in the presence of mixed substrates, including lactate/FeS and lactate/Fe(0) (13, 18). However, previous cultures did not grow on lactate alone (13, 18). Thus, while the growth in the presence of combined organic and inorganic electron donors suggests a potential for mixotrophy, further work is needed to clarify exactly which substrates promote *Sideroxydans* sp. CL21 growth and support organoheterotrophy.

Markers of iron oxidation activity could be the genes within iron oxidation pathways, but our knowledge of these pathways is still in development. The *Sideroxydans* sp. CL21 genome encodes several genes related to iron oxidation and associated pathways. Its genome encodes two known iron oxidases, 1) the monoheme cytochrome-porin Cyc2 and 2) the decaheme cytochrome MtoA and associated porin MtoB (17–19). In fact, the genome has three copies of *cyc2* and an *mtoAB,* plus an additional copy of *mtoA* next to an unnamed porin. While the presence of *cyc2* or *mtoA* indicate iron oxidation capability, it is unclear if they are suitable genetic markers of iron oxidation *activity*. Ideally, a gene marker would exhibit little to no expression when there is no iron oxidation activity and be upregulated during iron oxidation, with expression level scaling with activity. Yet, in the closely related *Sideroxydans lithotrophicus* ES-1, one copy of *cyc2* is highly expressed in both iron-oxidizing and thiosulfate-oxidizing cells, while the other two copies have low expression during thiosulfate oxidation, but are upregulated when oxidizing iron (10). In marine hydrothermal environments, the *cyc2* gene is highly expressed by various iron-oxidizing Zetaproteobacteria, and expression increases in microcosms amended with Fe(II) (20). In the facultative FeOB Zetaproteobacteria *Ghiorsea bivora* TAG-1, *cyc2*_TAG-1_1_ and Cyc2_TAG-1_1_ are expressed highly under both H_2_- and iron-oxidizing conditions (21). Together, this suggests *cyc2*/Cyc2 expression may not necessarily correspond to iron oxidation activity. There is less information on *mtoAB/*MtoAB, though these iron oxidase genes/proteins are upregulated in *S. lithotrophicus* ES-1 grown on Fe(II) minerals (22, 23). *Sideroxydans* sp. CL21 also encodes two gene clusters, predicted to encode porin-cytochrome complexes (PCC), each with two multiheme cytochromes (PCC3_copy1_, cytochromes have 24 and 14 hemes; PCC3_copy2_, cytochromes have 24 and 12 hemes). PCC3 is hypothesized to also play a role in extracellular electron transport and possibly iron oxidation (24). Additional putative components of the iron oxidation pathway (*mtoD*, *cymA/imoA*, Slit 1321-1324_ES-1_) are less well studied; these include genes for inner membrane and periplasmic electron carriers. In all, *Sideroxydans* sp. CL21 has a wide variety of genes that are known or predicted to participate in iron oxidation, giving multiple possibilities for marker genes.

Here, we constrain potential iron oxidation marker genes by studying differential gene expression in *Sideroxydans* sp. CL21. We first optimized the cultivation conditions by testing *Sideroxydans* sp. CL21’s growth on organic acids alone, plus other Fe(II) and non-iron substrates. Cells grew quickly and to the highest cell density on organic acids. We then tested the ability of these organic-grown cells to oxidize iron and found that lactate-grown cells exhibited a difference in iron-oxidizing phenotype between mid-log (no iron oxidation activity) and late-log (iron oxidation activity). This enabled us to compare gene expression between iron-oxidizing and non-iron-oxidizing conditions to detect genes whose expression corresponds to iron oxidation potential. The genes identified help expand our knowledge of the iron oxidation pathway and provide a list of candidate marker genes for iron oxidation.

## Results

### Physiology and growth on Fe(II), S, and organic substrates

*Sideroxydans* sp. CL21 grew on a range of individual organic and inorganic electron donors (Fig. 1). *Sideroxydans* sp. CL21 grew well on dissolved, ferrous iron (FeCl_2_), reaching a cell density of 1.25 × 10^7^ cells/mL in 15 days (Fig. 1A). This growth on FeCl_2_ was improved when media included 2.5 mM sodium citrate to complex iron, reaching a density of 2.76 × 10^7^ cells/mL (Fig. 1A). Attempts to culture *Sideroxydans* sp. CL21 on 5 mM sodium citrate alone showed no growth beyond that of a no-substrate control (Fig. 1A), demonstrating *Sideroxydans* sp. CL21 cannot conserve energy from citrate. As Fe(II) was the only available electron donor, these results confirm that *Sideroxydans* sp. CL21 can grow as a lithoautotrophic iron oxidizer.

**FIG 1.**
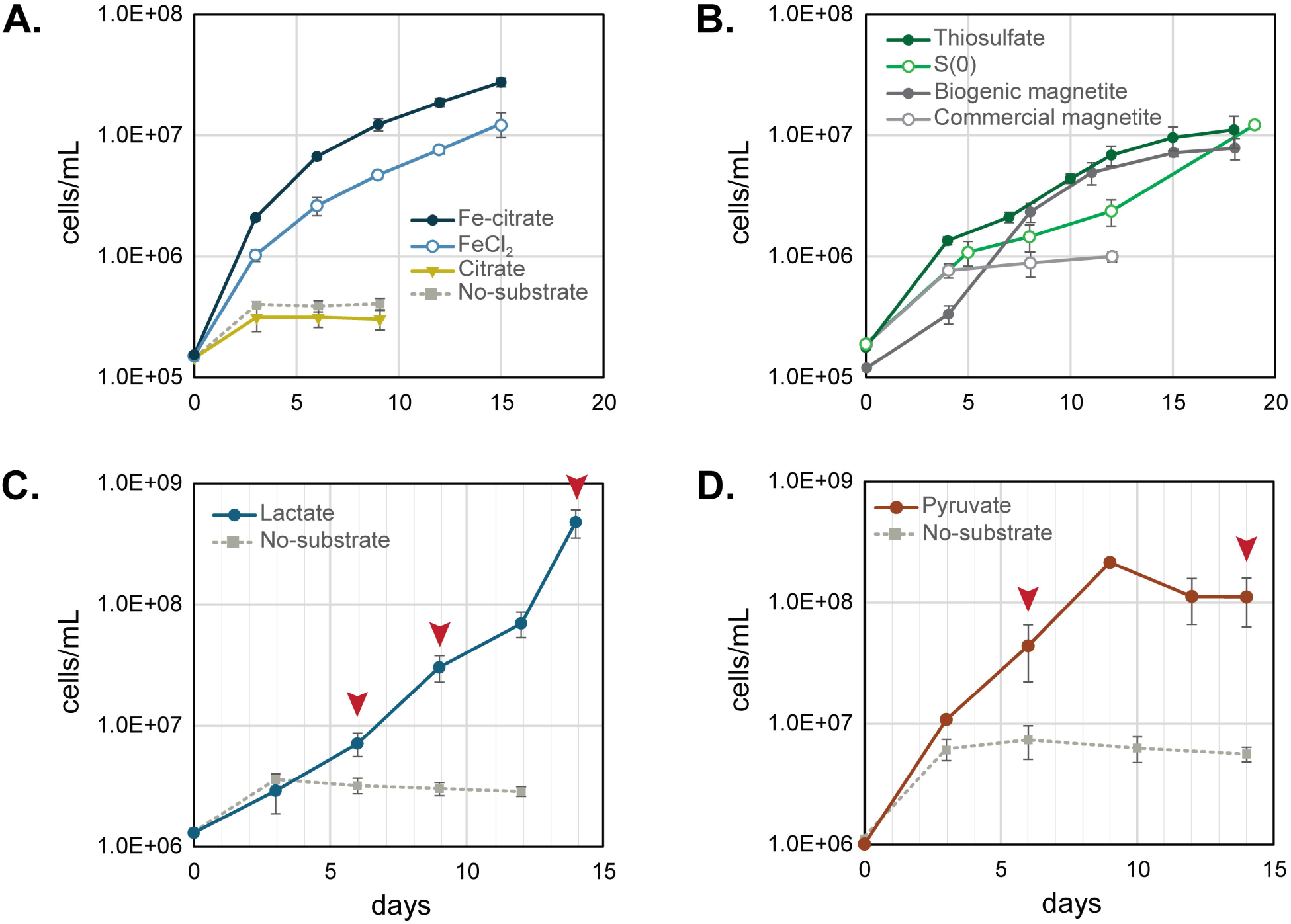
Growth of *Sideroxydans* sp. CL21 on: (A) FeCl_2_, FeCl_2_ + citrate, and citrate compared to a no substrate control; (B) thiosulfate, S(0), biogenic and commercial magnetite; (C) lactate, and (D) pyruvate. Red arrows in panels C indicate days cells were sampled for the Fe(II) spike experiments and transcriptome sequencing. Red arrows in panel D indicate days cells were sampled for Fe(II) spike experiments. Error bars represent one standard deviation between three replicates.

*Sideroxydans* sp. CL21 was cultured on commercial and biogenic magnetite to assess its growth on solid iron minerals. Cells grew better on the biogenic versus commercial magnetite (Fig. 1B). This may be due to the lower crystallinity and higher solubility of biogenic magnetite, which sheds more Fe(II) into solution than commercial magnetite (25). The biogenic magnetite, which was produced from ferrihydrite by the iron-reducer *Shewanella*, may also contain extracellular metabolites that could enhance iron oxidation activity and growth in *Sideroxydans* sp. CL21 (26).

*Sideroxydans* sp. CL21 was also tested for growth on sulfur substrates. The genome includes *dsrAB* genes that encode a reverse dissimilatory sulfite reductase (rDSR) (19). It has previously been cultured on FeS and in agarose-stabilized media in gradient tubes with either thiosulfate or thiosulfate plus H_2_ or lactate (18). Since agarose may provide a source of C (27, 28), we used liquid media without an organic carbon source to confirm whether cells could grow using individual sulfur substates as their sole electron donor. In these experiments *Sideroxydans* sp. CL21 grew on thiosulfate and elemental sulfur (Fig. 1B), reaching densities near 10^7^ cells/mL. However, growth was slower (18-19 days) and cell density was lower than that of Fe(II)-grown cells (Fig. 1A, 1B). Nonetheless, this shows *Sideroxydans* sp. CL21 is capable of two additional lithoautotrophic metabolisms using thiosulfate and S(0) as electron donors.

Previous studies showed growth of *Sideroxydans* sp. CL21 in multi-substrate experiments where lactate was combined with inorganic substrates (FeS, Fe(0), H_2_, NaS_2_O_3_) in agarose gradient tubes (18), yet cells were never successfully grown on lactate alone using these methods. Our data show that *Sideroxydans* sp. CL21 grew in liquid culture with either 5 mM lactate (Fig. 1C, 2A) or 5 mM pyruvate (Fig. 1D, 2B) as the sole electron donor. It is unclear why growth on lactate was positive in our study but previously negative, but it could be due to the more controlled oxygen concentrations in the headspace of these serum bottle liquid cultures versus a gradient tube. Cells cultured on lactate grew to an order of magnitude higher density than those cultured on Fe(II), reaching 4.8 × 10^8^ cells/mL in 14 days (Fig. 1C). Together, these results show *Sideroxydans* sp. CL21 can grow organotrophically, and likely organoheterotrophically, on lactate or pyruvate.

**FIG 2.**
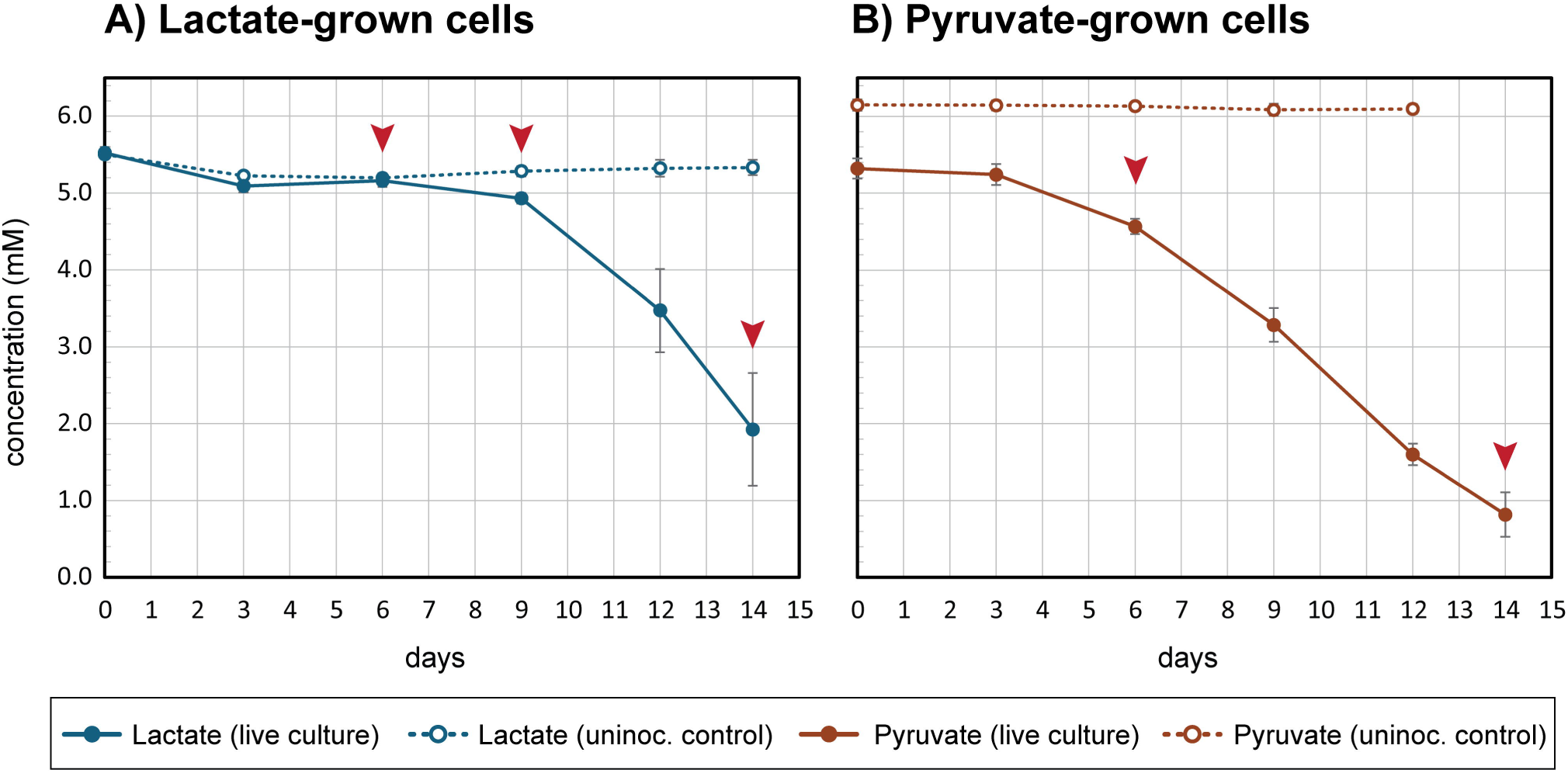
Consumption of (A) lactate and (B) pyruvate over time by live *Sideroxydans* sp. CL21 in growth experiments (Fig. 1C, 1D) versus uninoculated controls. Arrows indicate sampling points for short-term iron oxidation assays (Fig 3). Error bars represent one standard deviation between three replicates.

### The potential for mixotrophic energy metabolism

*Sideroxydans* sp. CL21 cells have the genomic potential to carry out multiple energy metabolisms simultaneously (mixotrophy), but this requires further experimental proof. Our approach was to determine whether *Sideroxydans* sp. CL21 cells grown on short chain organic acids could also oxidize ferrous iron in a short-term assay. We first harvested lactate-grown cells on days 6, 9, and 14 then added a 400 μM “spike” of FeCl_2_, and measured Fe^2+^ concentration over time. On Day 6, live cells (7.1 × 10^6^ cells/mL) did not oxidize iron within the 60-minute experiment. On Day 9 (3.3 × 10^7^ cells/mL), the modest amount of iron oxidation was mostly driven by a single bottle, 6A, which had a higher cell density than the other replicates, making it closer to late-log phase (Fig. 3A, Fig. S1). In contrast, with live cells (4.8 × 10^8^ cells/mL) harvested on Day 14, iron was completely oxidized within 15 minutes (Fig. 3A) despite the presence of >1 mM of lactate remaining in the cultures (Fig. 2A). This difference in response between mid- and late-log cultures indicates iron oxidation in *Sideroxydans* sp. CL21 is regulated. We also tested pyruvate-grown cells (Fig. 1D, 2B) and found *Sideroxydans* sp. CL21 was primed to oxidize iron on both Day 6 (4.4 × 10^7^ cells/mL) and Day 14 (1.1 × 10^8^ cells/mL) (Fig. 3B). Together, these results demonstrate that *Sideroxydans* sp. CL21 has the ability to oxidize iron while growing on organic acids, indicating that cells are capable of mixotrophic energy metabolism. Furthermore, because iron oxidation is regulated, we are able to use these samples to investigate gene expression differences between iron-oxidizing and non-iron-oxidizing conditions.

**FIG 3.**
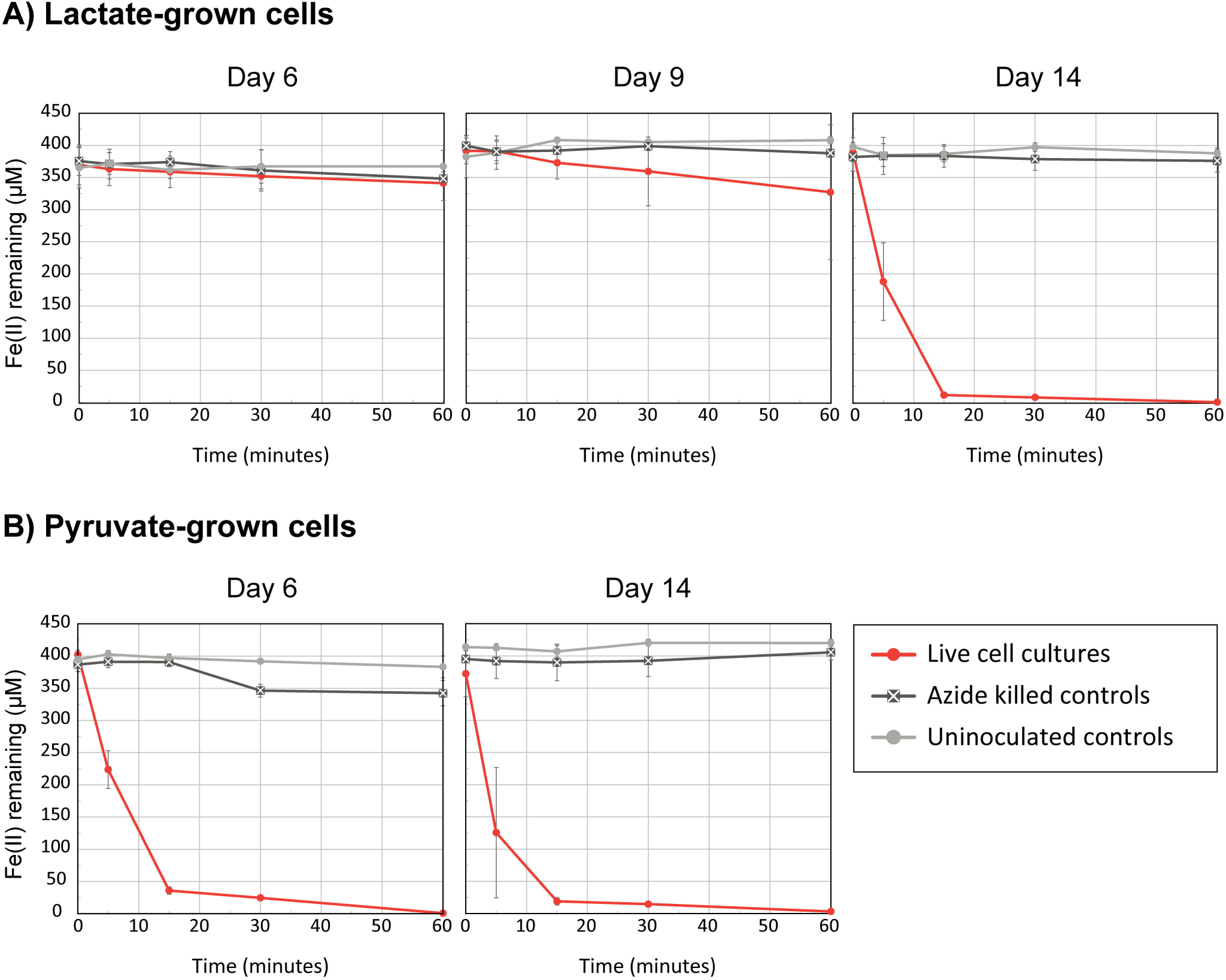
Results of Fe(II) spike assays for lactate- and pyruvate-grown cells. Aliquots of live, lactate-grown *Sideroxydans* sp. CL21 from the transcriptome experiment (Fig. 1C, 2A) rapidly oxidized iron on Day 14 compared to Day 6 or Day 9 (A). Cells from pyruvate growth experiments (Fig. 1D, 2B) rapidly oxidized iron on both Day 6 and Day 14 (B). Error bars represent one standard deviation between three replicates.

### Transcriptomics

Lactate-grown cultures were harvested for transcriptomics after aliquots were taken for the Fe(II)-spike experiments on Days 6, 9, and 14 (Fig. 1C, 2A). We present the transcriptome data as Z-scores, calculated from the normalized, regularized log (rlog) transformed read data. This gives a measure of a gene’s expression on a log scale relative to the mean expression (Z-score = 0) of all genes at a given time point.

#### Expression of lactate and carbon fixation genes

The *Sideroxydans* sp. CL21 genome contains six genes for lactate oxidation. These genes are co-located in the genome and encode a lactate response regulator (LldR), L-lactate dehydrogenase (LldE), L-lactate dehydrogenase (LldG), L-lactate dehydrogenase (LldF), D-lactate dehydrogenase (Dld), and lactate permease (LldP) (Fig. 4A). The genes *lldR*, *dld*, and *lldP* set *Sideroxydans* sp. CL21 apart from other closely related Gallionellaceae isolates *S. lithotrophicus ES-1* and *S. emersonii* MIZ01 that are unable to oxidize lactate (Fig. 4A). As expected, all six of the *Sideroxydans* sp. CL21 lactate oxidation genes were consistently expressed throughout the experiment (Fig. 4B, Table S1). None of the genes were significantly up- or down-regulated between the timepoints (Fig. 4B, Table S1), which fits with the consistent availability of lactate in the media (Fig. 2A).

**FIG 4.**
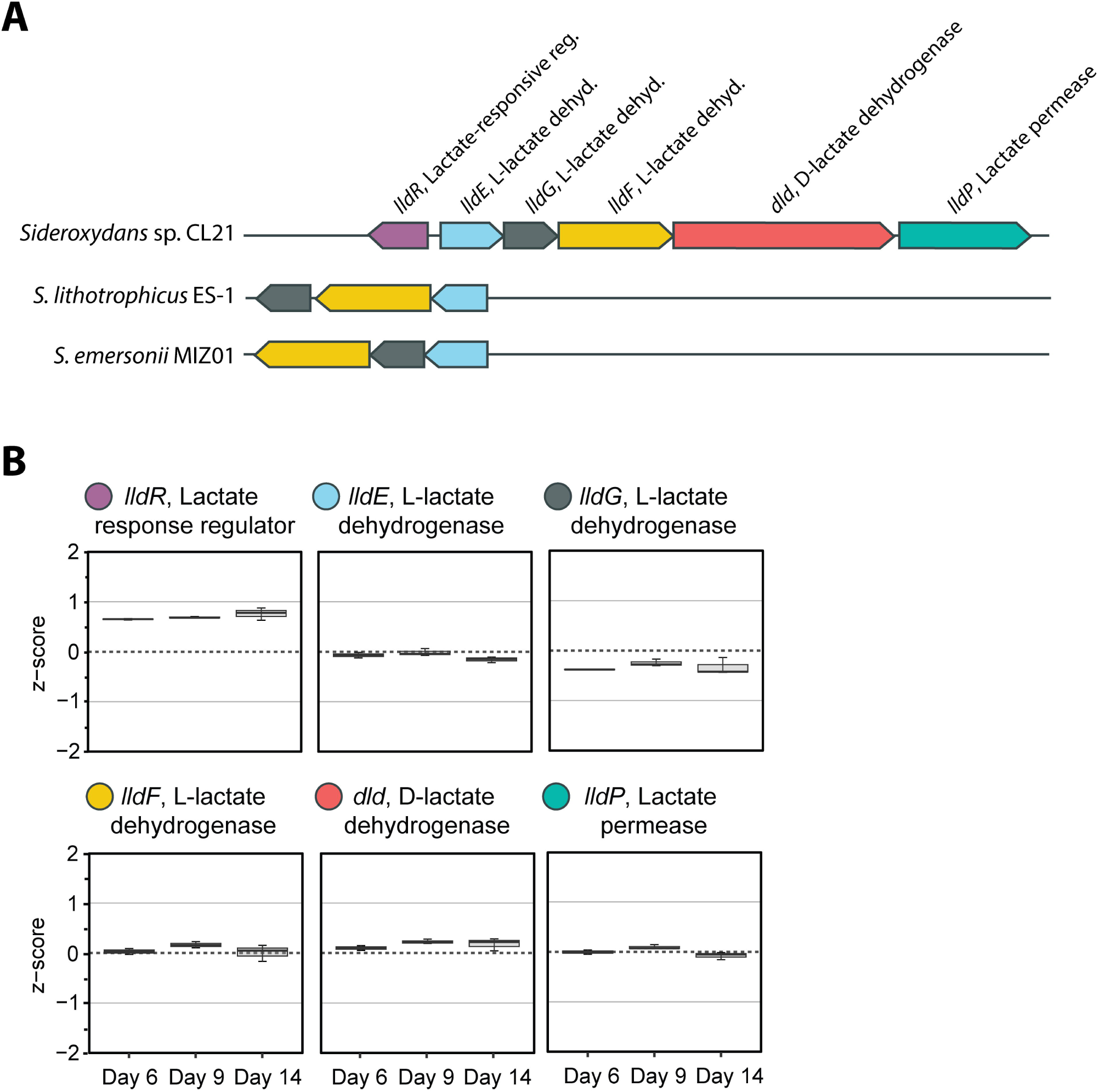
A) The six lactate oxidation genes encoded by *Sideroxydans* sp. CL21 and B) their patterns of expression on days 6, 9, and 14. Expression is represented by Z-score, which shows the expression of each gene relative to the mean gene expression (0) at the time point. Error bars represent the spread of the interquartile range.

When cultured with solely inorganic substrates, *Sideroxydans* sp. CL21 grows lithoautotrophically fixing CO_2_ via ribulose bisphosphate carboxylase (RuBisCo) and the Calvin–Benson–Bassham (CBB) cycle (26). High concentrations of organic carbon might be expected to promote heterotrophy and suppress autotrophy, yet lactate-grown *Sideroxydans* sp. CL21 still expressed genes for both Form I and Form II RuBisCo at all timepoints (Fig. S1B). Expression of Form I RuBisCo was relatively low while expression of Form II RuBisCo was high and greater than mean total expression at all three timepoints (Fig. S1B, Table S1). Form II RuBisCo has a higher affinity for oxygen (29), so its higher expression relative to Form I would be consistent with the micro-oxic culturing conditions used in our experiments. While further metabolic evidence is needed, high expression of the gene encoding Form II RuBisCo suggests *Sideroxydans* sp. CL21 could continue to fix CO_2_ while oxidizing lactate.

#### Expression of iron oxidation genes

The different iron oxidation activities of lactate cultures prompted us to investigate whether this ability corresponds to expression of iron oxidation genes. *Sideroxydans* sp. CL21 has multiple iron oxidase genes, three copies of *cyc2* and two copies of *mtoA*. Though there was variation in expression between the copies, overall both *cyc2* and *mtoA* were highly expressed on all three days (Fig. 5A, 5B). Compared to Day 6 (no iron oxidation), expression of *cyc2* or *mtoA* genes was not statistically significantly up- or down-regulated on Day 14, when iron oxidation was rapid (Fig. 5A, 5B, Table S1). Thus, expression of known iron oxidase genes did not correspond to cells’ ability to oxidize iron.

**FIG 5.**
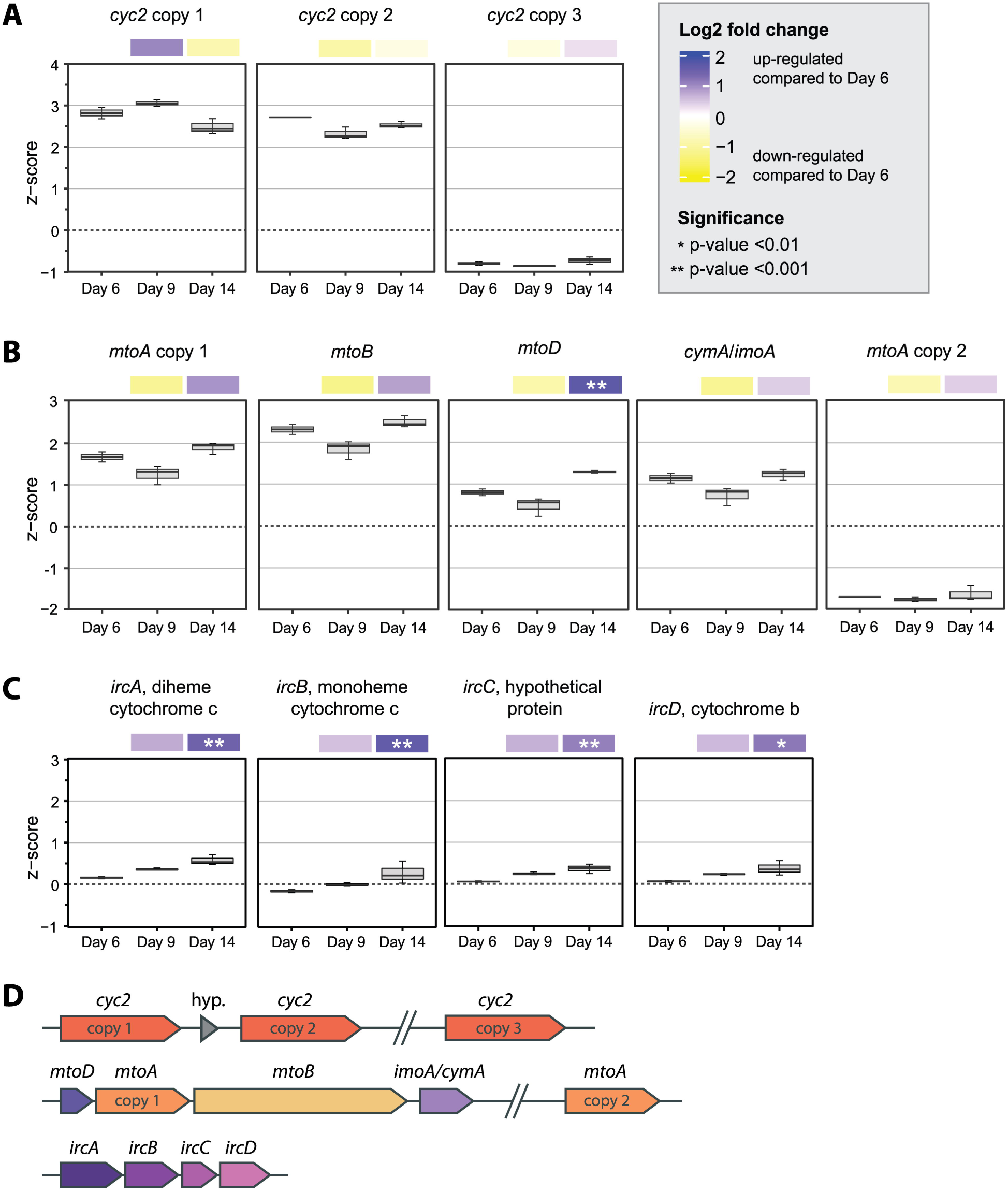
Boxplots of Z-score normalized expression for each time point with heatmaps showing the Log2 Fold Change in expression between Day 9 and Day 14 versus Day 6. Expression in the boxplots is represented by Z-score, which shows the expression of each gene relative to the mean gene expression (0) at the time point. Z-score was calculated from rlog normalized count data from DESeqs2. Heatmaps above the boxplots represent the Log2 Fold Change in average gene expression between Day 9 and Day 14 versus Day 6, with statistically significant p-values noted with asterisks (*). Data is shown for: (A) *cyc2* iron oxidase genes; (B) *mtoA* iron oxidase, *mtoB* porin, and *mtoD*, *imoA/cymA* periplasmic *c*-type cytochrome genes; and (C) *ircABCD* Fe(II)-responsive genes. Error bars represent the spread of the interquartile range. A gene map (D) shows the co-location of genes in the CL21 genome. Slashes breaking the line indicate genes are distant and “hyp.” denotes a gene encoding a hypothetical protein.

Both known iron oxidases, Cyc2 and MtoAB, are predicted to be outer membrane, porin-cytochromes, allowing them to oxidize Fe(II) outside the cell. Though functionally unverified, porin-cytochrome c complex 3 (PCC3) is predicted to be an outer membrane porin-cytochrome complex similar to, but larger than, the MtoAB iron oxidase complexes (24). PCC3 is widely distributed among various FeOB genomes (19, 24), making PCC3 another potential iron oxidation mechanism. *Sideroxydans* sp. CL21 has two copies of the PCC3 gene cluster. Each consists of four genes predicted to encode an extracellular multiheme cytochrome c (MHC), a porin, a periplasmic MHC, and an inner membrane protein. Analysis of the Z-scores, where mean expression is 0, showed both copies of PCC3 were expressed higher than the mean (Z-score ~1; Fig. S1), yet not as highly as *cyc2* or *mtoA* (Z-score ~3; Fig. 5A, 5B, Fig. S1) across all three time points. From Day 6 to Day 14, PCC3 copy 1 was downregulated while PCC3 copy 2 was upregulated, though the only significant (p-value <0.01) change was to the inner membrane protein of PCC3 copy 2 (Fig. 6, Fig. S1, Table S1).

**FIG 6.**
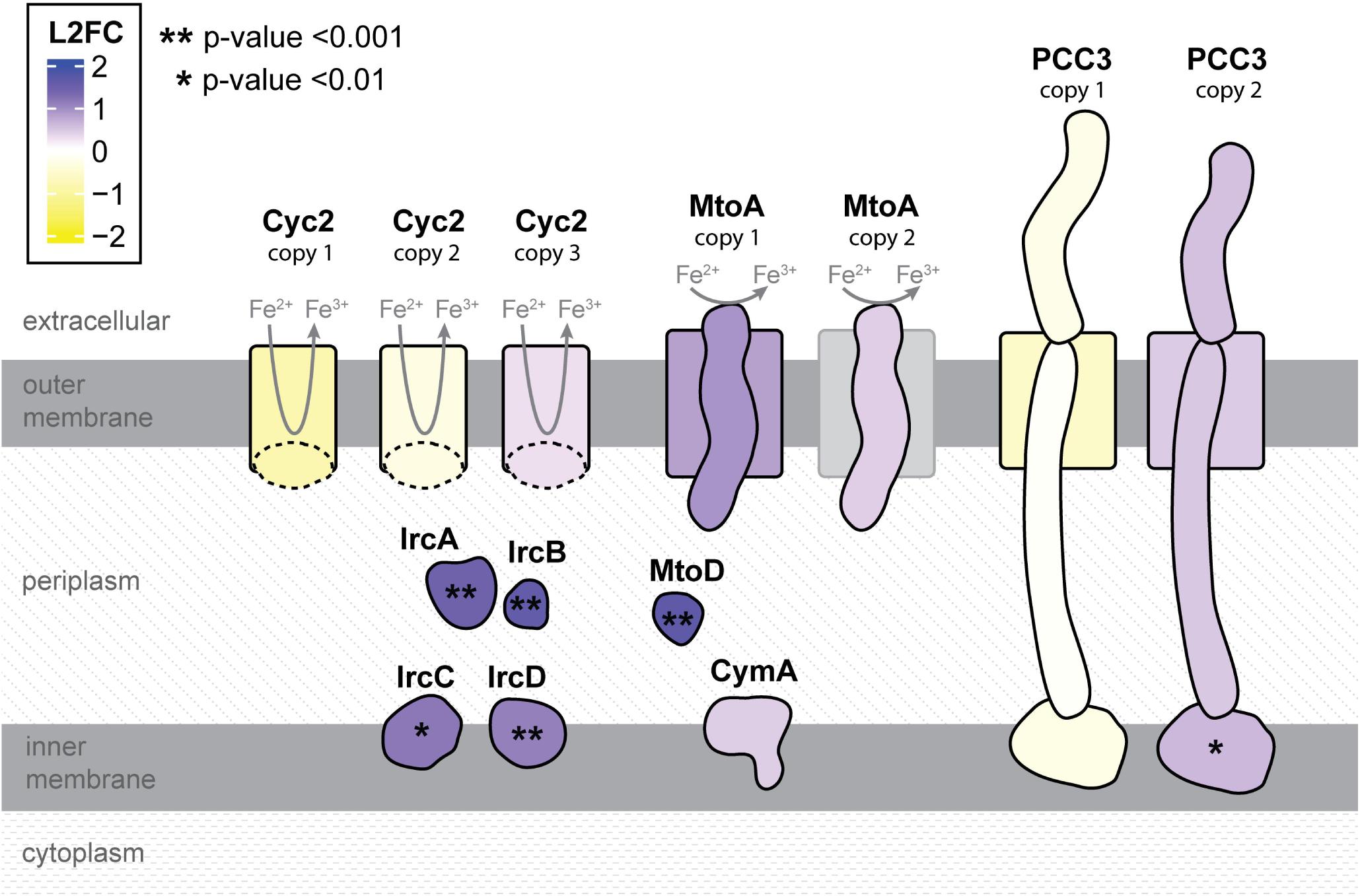
Cartoon schematic of iron oxidases, outer membrane cytochrome, porin, periplasmic, and inner membrane proteins colored by log2 fold change (L2FC) in normalized, rlog transformed gene expression between Day 14 and Day 6. Positive numbers (purple) were upregulated on Day 14 relative to Day 6. Asterisks indicate the gene encoding a particular protein was significantly upregulated.

Iron oxidases are just one part of FeOB extracellular electron transport pathways, which require additional cytochromes to convey electrons to the terminal oxidase or quinone pool. *Sideroxydans* sp. CL21 has genes encoding two *c*-type cytochromes, the monoheme MtoD and tetraheme CymA/ImoA, co-located in the genome next to *mtoA* copy 1 (Fig. 5D). Based on genomic context and previous genetic/biochemical characterization, these proteins may form part of the Mto iron oxidation pathway, moving electrons from the MtoA iron oxidase to the electron acceptor and/or quinone pool (30–33). Here, *mtoD* and *cymA/imoA* were upregulated at late log (Day 14) relative to Day 6, with *mtoD* being significantly upregulated (p-value <0.001) (Fig. 5, Fig. 6). In this case, the iron oxidation capability corresponds best with expression of the periplasmic electron carrier gene *mtoD*.

*Sideroxydans* sp. CL21 also has homologs to the four “Fe(II)-responsive” genes identified in *S. lithotrophicus* ES-1 (Slit_1321-Slit_1324 (Zhou et al. 2021)). We refer to these genes as “*ircABCD”* for “**<u>i</u>**ron **<u>r</u>**esponsive **<u>c</u>**luster.” The *irc* genes encode two *c*-type cytochromes, diheme (IrcA) and a monoheme (IrcB), both predicted to be periplasmic. They also encode a hypothetical protein (IrcC) and a *b*-type cytochrome (IrcD), with predicted localization in the inner membrane (Fig. 6). These genes were upregulated when *S. lithotrophicus* ES-1 was grown on Fe(II) versus thiosulfate (10) and also highly expressed in *Sideroxydans* sp. CL21 when grown with Fe(0) (26). In our lactate experiment, expression of these genes increased as cells approached late-log phase (Day 14) and all were significantly upregulated on Day 14 relative to Day 6 (Fig. 5C, Fig. 6). This significant upregulation corresponds to cells’ ability to oxidize iron on Day 14.

Overall, *Sideroxydans* sp. CL21 seems to be concurrently expressing genes for its known and putative iron oxidases (*cyc2*, *mtoA*, and PCC3) regardless of whether cells are capable of iron oxidation. However, expression of genes for periplasmic and inner membrane proteins (*mtoD*, *ircABCD*, inner membrane component of PCC3 copy 2) varies. These genes were significantly upregulated in cells that showed iron oxidation activity, making them potential markers of iron oxidation activity.

## Discussion

In this study, we demonstrated that *Gallionellaceae Sideroxydans* sp. CL21 exhibits remarkable metabolic versatility. *Sideroxydans* sp. CL21 can utilize a wide range of substrates, including short-chain organic acids such as lactate and pyruvate, as well as oxidize various soluble inorganic substrates, such as Fe(II) and thiosulfate. Additionally, it can metabolize mineral substrates, including magnetite and elemental sulfur (S(0)). Overall, our findings show that *Sideroxydans* sp. CL21 is capable of organotrophy and lithoautotrophy, with the potential for mixotrophy and organoheterotrophy. This metabolic flexibility likely enhances its competitiveness and enables it to contribute to a wide range of biogeochemical transformations. Understanding the ecophysiology and biogeochemical roles of facultative iron oxidizers like *Sideroxydans* sp. CL21 requires more than just detecting their presence in environmental samples. We need a means to decipher whether these FeOB are actively oxidizing iron. Developing methods to directly track microbial iron oxidation activity requires (1) the identification of genes encoding the iron oxidation pathway and (2) determining which of these genes can serve as reliable markers of microbial iron-oxidizing activity.

To be a marker of activity, a gene needs to increase in expression with iron oxidation and have minimal to no expression when cells are unable to oxidize iron. Validating marker genes requires study of facultative FeOB, of which there are few in culture, and there ideally needs to be a condition under which cells *cannot* oxidize iron. Previous studies on *S. lithotrophicus* ES-1 showed differential gene expression of cells grown on Fe(II)-citrate and thiosulfate, but thiosulfate-grown cells were able to oxidize iron quickly with no lag after the substrate was switched to Fe(II). In contrast, *Sideroxydans* sp. CL21 grown on lactate was surprisingly unable to oxidize iron until late-log phase (Day 14), giving an opportunity to compare gene expression of iron-oxidizing and non-iron-oxidizing conditions.

### Relationship between iron oxidation gene expression and activity

Although significant progress has been made in assigning function to the key iron oxidases such as Cyc2 and MtoA (10, 20, 33–35), the pathway is likely more complex, and additional iron oxidases may yet be discovered. Genes encoding the larger multiheme cytochrome complexes PCC3_1 and PCC3_2 are expressed at higher than mean levels. PCC3s are predicted to be large outer membrane cytochrome (OMC)-porin complexes (Fig. 6) (24), and while their function has not been tested, as OMCs, they could in principle oxidize iron. Thus, based on the *Sideroxydans* sp. CL21 transcriptome, there are at least four possible iron oxidases (Cyc2, MtoA, and the PCC3_1 and PCC3_2 cytochromes) with various proteins that could direct electrons to terminal oxidases or reverse electron transport pathways.

One might expect the expression of iron oxidase genes to correlate with iron oxidation, but our results show that the well-characterized iron oxidase genes *cyc2* and *mtoA* are always highly expressed in our experiments, and thus do not correspond to iron oxidation ability. This differs from *S. lithotrophicus* ES-1, which has very low expression of *mtoA* on dissolved Fe^2+^ but upregulates *mtoA* when grown on solid Fe(II)-containing minerals smectite and magnetite (10, 22, 23). We showed that *Sideroxydans* sp. CL21 grows on magnetite, and though magnetite was not present in lactate-grown cultures, expression of *mtoA* may still indicate a readiness to take up electrons from solid minerals. *Sideroxydans* sp. CL21 is more like *S. lithotrophicus* ES-1 in its high expression of *cyc2*. In *S. lithotrophicus* ES-1, expression of *cyc2* (one of three copies) is high even when cells are cultured on thiosulfate (10). Similarly, *Sideroxydans* sp. CL21 highly expresses two of its three *cyc2* copies when grown on lactate. Expression of *cyc2* is also generally high in marine iron-oxidizing Zetaproteobacteria (20). However, in *S. lithotrophicus* ES-1 additional copies of *cyc2* are iron-responsive (10). In theory, Cyc2 production could be post-transcriptionally regulated. However, this is not the case in *S. lithotrophicus* ES-1, since proteome data shows Cyc2 is highly expressed on Fe(II) (dissolved Fe^2+^ and magnetite (23)) and thiosulfate (25). One explanation for the overall high expression is that these FeOB maintain readiness or, in the case of *Sideroxydans* sp. CL21, partial readiness to oxidize iron. In any case, because *cyc2* expression does not correspond to iron oxidation activity, *cyc2* (alone) is not an accurate indicator of biotic iron oxidation.

Beyond iron oxidases, iron oxidation activity also requires electron carrier proteins in the periplasm and inner membrane. Currently, there is limited evidence supporting other essential components of the pathway, particularly periplasmic electron carriers and inner membrane proteins (Table S3). Our findings contribute to filling this gap by providing evidence for the involvement of genes encoding predicted periplasmic and inner membrane proteins. In fact, the genes that were most upregulated in cells that showed iron oxidation (Day 14) were periplasmic *c*-type cytochrome genes *ircA*, *ircB*, and *mtoD* (Fig. 6), followed by the inner membrane protein genes *ircC*, *ircD*, and the PCC3_2 inner membrane component. There is accumulating evidence for the involvement of *irc* genes in iron oxidation. Homologs are widespread among Gallionellaceae and found in several other FeOB including *Leptothrix cholodnii* SP-6, *Rhodoferax* spp., and several Zetaproteobacteria (Table S2; (19, 21)). In *S. lithotrophicus* ES-1, *irc* genes are upregulated on iron-oxidizing cultures versus thiosulfate-oxidizing ones (10). The periplasmic di- and mono-heme *c*-type cytochrome genes (*ircA* and *ircB*) are homologs of *grcJ* and *grcI* in *Ghiorsea bivoria* TAG-1. In *G. bivoria* TAG-1, GrcJ is significantly upregulated under iron-oxidizing vs. H_2_-oxidizing conditions and expression of *grcJ* increases over time when cells were grown with Fe(II) (21). In lactate-grown *Sideroxydans* sp. CL21, upregulated expression of *ircABCD* corresponded to the cells’ potential to oxidize iron in late-log cultures (Day 14). Thus, it appears that cells increase expression of electron carrier genes like *irc*, *grc*, and *mtoD* when cells are ready to oxidize iron, making them the best candidates thus far for markers of iron oxidation potential.

The results to date suggest that the iron oxidation pathway is not regulated as a complete set in the neutrophilic FeOB Gallionellaceae. Instead, at least one OMC (one Cyc2) is constitutively expressed, which could serve multiple purposes. Cyc2 may operate as an Fe(II) sensor to tell the cell when it is time to increase expression of the iron oxidation pathway. An issue with this pathway is that OMCs are complicated to produce; they must be secreted to the periplasm, where the heme cofactor is attached and then inserted into the outer membrane with proper folding. Each of these three steps involves separate machinery, so this requires substantial space in the inner membrane and periplasm, which may interfere with the electron transfer pathway. Thus, expressing Cyc2 and other OMCs ahead of iron oxidation may be a strategy for a quicker iron oxidation response. In contrast to the OMC, many of the periplasmic and inner membrane components are smaller and would be easier to produce when they are required. This strategy also allows the organism to express different periplasmic and inner membrane proteins to match different iron substrates (Fe(II), magnetite, chelated Fe(II)), which have varied redox potentials (36, 37), as is the case in *Geobacter* (38).

Thus, we conclude that the Gallionellaceae may implement a multistage production of the iron oxidation pathway, with a subset of the OMCs expressed proactively, followed by additional OMCs and the rest of the pathway once iron oxidation is required. This may be further tested as additional facultative FeOB are isolated. In the meantime, we will need to carefully parse the expression profiles to interpret FeOB activity in the environment, though the *irc*/*grc* genes are promising markers.

### C utilization among FeOB

*Sideroxydans* sp. CL21 is distinct among the current Gallionellaceae isolates in its ability to grow on organic acids. However, other uncultured mixotrophic or organo(hetero)trophic FeOB are likely present in C-rich environments. A pangenomic analysis of *Gallionellaceae* isolates and environmental MAGs reveals that the potential for lactate utilization is rare but not unique, as it has been observed in some genomes from the Crystal Bog humic lake (Wisconsin) and freshwater lakes near Kuujjuarapik-Whapmagoostui (Canada) (19, 39–41). Active organic substrate utilization by FeOB fundamentally alters their role in the carbon cycle. While FeOB are traditionally considered autotrophs that fix CO, the ability to grow organotrophically suggests they may also contribute to the degradation of organic carbon in situ. The ability to grow on organics may provide a significant ecological advantage in specific habitats, especially wetlands and peatlands. In peatlands, mixotrophic FeOB face competition with both aerobic and anaerobic microorganisms for organic intermediates such as lactate and pyruvate. However, FeOB exhibit a unique advantage by coupling the consumption of these substrates to aerobic respiration under microaerophilic conditions. This metabolic flexibility of FeOB enables them to thrive in environments like peatlands, where fluctuating water tables create dynamic oxic-anoxic gradients, offering transient microaerophilic niches. Thus, such versatility may be key to their survival and dominance in specific ecosystems, and play integral roles in biogeochemical iron and C cycling in dynamic environments.

### Implications of FeOB organotrophy for wetland C cycling

By demonstrating the versatility of FeOB metabolism and its genetic basis, our findings provide a foundation for understanding the broader biogeochemical roles of FeOB in wetland ecosystems and how they connect iron and C cycles. It is already recognized that organotrophic iron-reducing bacteria can compete for organic carbon, thus shunting carbon away from methanotrophs, which could reduce wetland methane emissions (42–46). FeOB help promote this process by replenishing the supply of Fe(III). If the FeOB are autotrophic, then iron oxidation drives the regeneration of organic C, partly reversing the effect of organotrophic iron reduction and muting the effects on methanogenesis. But if the FeOB are organotrophic, they would join the competition for organics and, if sufficiently abundant and active, contribute to the suppression of methanogenesis. These are theoretical effects; evaluating the actual effects of FeOB organotrophy requires specifically monitoring FeOB C metabolism in the wetland environment and soil incubations. The potential iron oxidation gene markers would give us a way to link iron oxidation activity to organotrophic metabolisms via genome-resolved metatranscriptomics, paving the way toward deeper insights into FeOB influences on wetland biogeochemistry and interactions with wetland communities. This knowledge is essential for predicting how iron and C cycling—and associated greenhouse gas fluxes—will respond to environmental changes.

## Methods

### Growth experiments

For all growth experiments, *Sideroxydans* sp. CL21 was cultured in triplicate, in 150 mL serum bottles with 80 mL of modified Wolfe’s minimal medium (MWMM; containing 1.0 g NH_4_Cl, 0.5 g MgSO_4_⋅7H2O, 0.2 g CaCl_2_, and 0.05 g K_2_HPO_4_ per liter). All cultures were amended with a 0.1% vol/vol addition each of ATCC trace vitamins and trace minerals. To keep micromolar concentrations of oxygen, headspaces were flushed daily with low-oxygen gas mix (2% O_2_, 20% CO_2_, 76% N_2_). A Firesting oxygen spot sensor (Pyro Science) was used to measure and monitor oxygen concentrations. With the exception of lactate and pyruvate cultures, all bottles were incubated on their sides in a dark culturing cabinet at room temperature. Lactate and pyruvate bottles were incubated upright in a New Brunswick Innova 42 shaking incubator at 20°C, 75 RPM. Growth for all cultures was measured by direct cell counting whereby cells were stained with Syto-13 then counted manually in a Petroff-Hauser counting chamber under fluorescence microscopy.

Lactate cultures included a one-time addition of sodium DL-lactate (CAS 72-17-3) for a concentration of 5 mM and were buffered with 10mM of 2-(N-morpholino)ethanesulfonic acid (MES) at pH 6.0. Likewise, citrate (CAS 6132-04-3), pyruvate (CAS 113-24-6), thiosulfate (CAS 10102-17-7), and S(0) cultures were buffered with MES at pH 6.0 and given a one-time dose of sodium citrate dihydrate, sodium pyruvate, sodium thiosulfate pentahydrate, or elemental sulfur powder for a starting concentration of 5 mM. Biogenic and commercial magnetite cultures were each given a single addition of magnetite (1 g/L). Biogenic magnetite was synthesized by incubating *Shewanella oneidensis* MR-1 in Luria-Bertani (LB) media with two-line ferrihydrite for 10-12 days in an anerobic chamber. After incubation a magnet was used to hold the magnetite in place while the supernatant was decanted. Minerals were washed 3x with deoxygenated water, then dried in an anaerobic desiccator.

FeCl_2_ cultures were buffered with 50 mM of MES at pH 6.0. These cultures were given a daily addition of FeCl_2_ ranging from 250 μM on Day 0 to 500 μM on Day 14. The FeCl_2_ + citrate experiment was conducted under these same conditions plus a one-time addition of 2.5 mM sodium citrate.

### Organic acids measurements

Samples from lactate and pyruvate cultures were collected, filtered with a 0.22 μm syringe filter, and stored at 4°C. Concentration of organic acids (lactate, pyruvate) were measured by high-performance liquid chromatography (HPLC). Standards for formate and acetate were also included to check for partial oxidation products, but neither were detected. Measurements were made on a Shimadzu HPLC with a Photodiode Array Detector (model MPD-40) with a Alltech Prevail Organic Acid 5 µm 150 × 4.6 mM column and guard column. Organic acids were eluted at room temperature with 25 mM KH_2_PO_4_ at pH 2.5, adjusted with phosphoric acid, at 1 mL/minute. Absorbance was read at 210 nm to quantify compounds based on peak area. Instrument control and data collection were performed using LabSolutions software (v. 5.106 SP1). Peak areas were exported to Microsoft Excel where calibration curves were generated by linear regression of authentic standard peak areas and used to calculate concentrations.

### Transcriptome experiment

*Sideroxydans sp.* CL21 was cultured in 5 mM lactate as described above. Bottles were inoculated with stationary phase, lactate-adapted cells. Cell counts and lactate measurements were taken on days 0, 3, 6, 9, 14. On days 6, 9, and 14, aliquots of cultures were used in Fe(II) spike assays (described below) and bottles were sacrificed in biological sextuplets for transcriptome sequencing. For all sacrifices, a 1:10 vol/vol of stop solution (buffer-saturated phenol:absolute ethanol at a 1:9 vol/vol) was injected into the bottles to stop transcription. Cells were then harvested by filtering onto a 0.22 μm polyethersulfone (PES) filter and stored at −80°C until RNA extraction.

### Fe(II) spike assay

Fe(II) spike assays were performed at different timepoints to test *Sideroxydans* sp. CL21’s response to Fe(II) when grown on lactate and pyruvate. For these assays, 10 mL of culture were extracted from each of the six inoculated serum bottles. Of this, 5 mL was incubated in 5 mM sodium azide for 20 minutes to create the azide-killed control. The other 5 mL was untreated and served as the “live cell” samples. A 5 mL aliquot was taken from six uninoculated bottles for the uninoculated control. All samples were bubbled with a 2% O_2_, 20% CO_2_, 78% N_2_ gas mix throughout the experiment to keep oxygen levels consistent and low. Each sample, live and control, was spiked with 20 μL of 100 mM FeCl_2_ for a 400 μM starting concentration. Samples were collected at 0, 5, 15, 30, and 60 minutes and preserved in a 1:1 mix with 40 mM sulfamic acid. After all samples were collected, a spectrophotometric ferrozine assay was used to measure ferrous iron concentrations (47, 48).

### RNA extraction and sequencing

RNA was extracted from cells collected on PES filters using a RNeasy PowerSoil RNA kit (Qiagen). Extracted RNA was treated with three RNase inhibitors (1 μL Invitrogen SUPERase, 3 μL RNaseOUT, 1.25 μL Ambion RNase Inhibitor) to prevent degradation. Total RNA was quantified using a Qubit RNA HS assay kit from Invitrogen and quality was assessed with an Agilent fragment analyzer. A Zymo RNA Clean & Concentrator™-5 kit was used to remove excess salts and DNA from all samples. After clean-up, three samples with the best yield and quality from each timepoint (Day 6 - 12.6, 13.2, 29.6 ng/µL; Day 9 – 34.6, 43.0, 25.6 ng/µL; Day 14 – 82.0, 20.0, 92.0 ng/µL), along with sample 6A from Day 9 (33.4 ng/µL) were selected for sequencing. Ribosomal RNA depletion and library preparation were done using a Zymo-Seq RiboFree^®^ Total RNA Library Kit. Sixty-four ng of total RNA from each samples was sequenced on an Illumina NextSeq 2000 using a P2 100-cycle protocol yielding up to 40 GB of data (up to 400 million single-end reads). All clean-up, library prep and sequencing was performed by the University of Delaware DNA Sequencing & Genotyping Center.

### Transcriptome QC and analysis

Raw transcriptome reads were quality checked using FastQC v0.11.9 (49). Reads were trimmed and adaptor sequences were removed using TrimGalore! v.0.6.6 (50) and Cutadapt v4.4 (51). A minimum quality score of 28 and minimum length of 75 bp were applied to filter out poor reads. Post-trim, the read quality was reassessed with FastQC v0.11.9 (49) and SortMeRNA v4.3.6 (52) was run on all samples to remove any rRNA sequences before mapping.

Reads were mapped to the closed, RefSeq annotated genome of *Sideroxydans* sp. CL21 (GCF_902459525.1) using Bowtie 2 v2.5.1 (53) and Samtools v1.10 (54). The average alignment rate per sample was 97%, with the exception of Day 6, Sample 3B which was 73%. Read counts were reported using Htseq v2.0.2 (55). After comparing the total number of reads mapped for each sample and the overall alignment rates, we determined that Day 6, Sample 3B was an outlier with potential contamination and too few quality reads to be useful. To be thorough, we ran and compared the DGE analysis (described below) with and without Day 6 Sample 3B. Exclusion of Day 6 Sample 3B did not affect the overall findings or change whether the differential expression of our genes of interest was significant. Instead, removing the outlier decreased noise and made the dataset more precise.

Differential gene expression was analyzed in R v4.4.0 (56) with RStudio v2023.06.0 Build 421 (57) using the DESeq2 v1.26.0 (58) package. Data was normalized using DESeqs default algorithm which uses an estimation of size factors to control for differences in sequencing depth. P-values were determined with the Wald Test and adjusted for false discovery rate (FDR) using the Benjamini-Hochberg (BH) correction. An alpha of 0.01 was set as the cut-off for significance. Reports of normalized count data for each sample were generated using the regularized logarithm (rlog) transformation (58). This rlog counts data was used to calculate Z-score to analyze the variance of individual genes across samples. To test validity of these transformations, we checked the expression of housekeeping genes for DNA gyrase and RNA polymerase *(gyrA, gyrB*, and *rpoA*). Their even expression over all timepoints indicates the rlog and Z-score transformations were successful (Fig. S1A, Table S1).

### Identification and analysis of the Fe(II)-responsive (*irc*) gene cluster

Homologs of the Fe(II)-responsive (*irc*) gene cluster were identified by using cblaster (59) to search the NCBI database for BLAST hits (e-value cut-off < 1E-10) to the *irc* genes of *Sideroxydans* sp. CL21 that were located within 20,000 bp of each other in their respective genomes. Positive hits needed to contain at least three of the genes, excluding the Hsp33 chaperone. Results were visualized using clinker (60). PSORTb 3.0 (61), Gneg-mPLoc (62), PredictProtein (63), and InterProScan (64, 65) were used to predict subcellular localization and domains of the *irc* genes.

### Data availability

Transcriptomic data were uploaded to NCBI BioProject PRJNA1216980.

## Acknowledgements

This work was funded by the National Science Foundation (EAR-1833525 to C.S.C., MCB-1817651 to C.S.C.) and the Office of Naval Research (N00014-17-1-2640 C.S.C.). R.L.H. was also supported by fellowships from University of Delaware Graduate College and the Microbiology Program/Unidel Foundation. Support from the University of Delaware Center for Bioinformatics and Computational Biology Core Facility (RRID:SCR_017696), including the use of the BIOMIX computer cluster, was made possible through funding from Delaware INBRE (NIH P20GM103446), the State of Delaware, and the Delaware Biotechnology Institute. KK was supported by the Collaborative Research Centre Chemical Mediators in Complex Biosystems (SFB 1127 ChemBioSys, Project Number 239748522), which is funded by the Deutsche Forschungsgemeinschaft.

We thank Thomas Hanson for his help with the HPLC analyses, Christian Sinjari for his assistance culturing *Sideroxydans* sp. CL21 on magnetite, Jessica Keffer for feedback and editing, and Mark Shaw for sample clean-up and sequencing.

We declare no conflict of interest.

## Supplementals

### Figures

- **Figure S1** – Line graphs of iron oxidation response of individual bottles on A) Day 6, B) Day 9, and C) Day 14.
- **Figure S2** – Boxplot/heatmap of A) housekeeping genes, B) RuBisCo, C) PCC3 copy 1, D) PCC3 copy 2

### Tables

- **Table S1** – Differential expression, normalized expression, and Z-score data for key genes
- **Table S2** – *ircABCD* homologs in FeOB and FeRB
- **Table S3** – Table of existing evidence for the involvement of various genes and proteins in iron oxidation and/or extracellular electron transport.

## Notes

### Competing Interest Statement

The authors have declared no competing interest.

### Summary of Updates

Minor edits were made in response to journal reviewer comments. Gene maps were added to Figure 5.

## References

1. Borch T, Kretzschmar R, Kappler A, Cappellen PV, Ginder-Vogel M, Voegelin A, Campbell K. 2010. Biogeochemical redox processes and their impact on contaminant dynamics. Environ Sci Technol 44:15–23.

2. Barros MEC, Rawlings DE, Woods DR. 1984. Mixotrophic growth of a *Thiobacillus ferrooxidans* strain. Appl Environ Microbiol 47:593–595.

3. Akob DM, Hallenbeck M, Beulig F, Fabisch M, Küsel K, Keffer JL, Woyke T, Shapiro N, Lapidus A, Klenk H-P, Chan CS. 2020. Mixotrophic iron-oxidizing *Thiomonas* isolates from an acid mine drainage-affected creek. Appl Environ Microbiol 86:e01424–20.

4. Fleming EJ, Woyke T, Donatello RA, Kuypers MMM, Sczyrba A, Littmann S, Emerson D. 2018. Insights into the fundamental physiology of the uncultured Fe-oxidizing bacterium *Leptothrix ochracea*. Appl Environ Microbiol 84:e02239–17.

5. Hallbeck L, Pedersen K. 1991. Autotrophic and mixotrophic growth of *Gallionella ferruginea*. Microbiology, 137:2657–2661.

6. Tothero GK, Hoover RL, Farag IF, Kaplan DI, Weisenhorn P, Emerson D, Chan CS. 2024. *Leptothrix ochracea* genomes reveal potential for mixotrophic growth on Fe(II) and organic carbon. Appl Environ Microbiol 0:e00599–24.

7. Sobolev D, Roden EE. 2004. Characterization of a neutrophilic, chemolithoautotrophic Fe(II)-oxidizing β-Proteobacterium from freshwater wetland sediments. Geomicrobiol J 21:1–10.

8. Eggerichs T, Wiegand M, Neumann K, Opel O, Thronicker O, Szewzyk U. 2020. Growth of iron-oxidizing bacteria *Gallionella ferruginea* and *Leptothrix cholodnii* in oligotrophic environments: Ca, Mg, and C as limiting factors and G. ferruginea necromass as C-source. Geomicrobiol J 37:190–199.

9. Emerson D, Moyer C. 1997. Isolation and Characterization of novel iron-oxidizing bacteria that grow at circumneutral pH. Appl Environ Microbiol 63:9.

10. Zhou N, Keffer JL, Polson SW, Chan CS. 2021. Unraveling Fe(II)-oxidizing mechanisms in a facultative Fe(II) oxidizer, *Sideroxydans lithotrophicus* strain ES-1, via culturing, transcriptomics, and reverse transcription-quantitative PCR. Appl Environ Microbiol 88:e01595–21.

11. Kato S, Itoh T, Iino T, Ohkuma M. 2022. *Sideroxyarcus emersonii* gen. nov. sp. nov., a neutrophilic, microaerobic iron- and thiosulfate-oxidizing bacterium isolated from iron-rich wetland sediment. Int J Syst Evol Microbiol 72.

12. Garry M, Farasin J, Drevillon L, Quaiser A, Bouchez C, Le Borgne T, Coffinet S, Dufresne A. 2024. *Ferriphaselus amnicola* strain GF-20, a new iron: and thiosulfate-oxidizing bacterium isolated from a hard rock aquifer. FEMS Microbiol Ecol 100:fiae047.

13. Lüdecke C, Reiche M, Eusterhues K, Nietzsche S, Küsel K. 2010. Acid-tolerant microaerophilic Fe(II)-oxidizing bacteria promote Fe(III)-accumulation in a fen. Environ Microbiol 12:2814–2825.

14. Küsel K, Blöthe M, Schulz D, Reiche M, Drake HL. 2008. Microbial reduction of iron and porewater biogeochemistry in acidic peatlands. Biogeosciences 5:1537–1549.

15. Hädrich A, Taillefert M, Akob DM, Cooper RE, Litzba U, Wagner FE, Nietzsche S, Ciobota V, Rösch P, Popp J, Küsel K. 2019. Microbial Fe(II) oxidation by *Sideroxydans lithotrophicus* ES-1 in the presence of Schlöppnerbrunnen fen-derived humic acids. FEMS Microbiol Ecol 95:fiz034.

16. Kügler S, Cooper RE, Wegner C-E, Mohr JF, Wichard T, Küsel K. 2019. Iron-organic matter complexes accelerate microbial iron cycling in an iron-rich fen. Sci Total Environ 646:972–988.

17. Cooper RE, Wegner C-E, McAllister SM, Shevchenko O, Chan CS, Küsel K. 2020. Draft genome sequence of *Sideroxydans* sp. strain CL21, an Fe(II)-oxidizing bacterium. Microbiol Resour Announc 9:e01444–19.

18. Cooper RE, Finck J, Chan C, Küsel K. 2023. Mixotrophy broadens the ecological niche range of the iron oxidizer *Sideroxydans* sp. CL21 isolated from an iron-rich peatland. FEMS Microbiol Ecol 99:fiac156.

19. Hoover RL, Keffer JL, Polson SW, Chan CS. 2023. Gallionellaceae pangenomic analysis reveals insight into phylogeny, metabolic flexibility, and iron oxidation mechanisms. mSystems 8:e00038–23.

20. McAllister SM, Polson SW, Butterfield DA, Glazer BT, Sylvan JB, Chan CS. 2020. Validating the Cyc2 Neutrophilic Iron oxidation pathway using meta-omics of Zetaproteobacteria iron mats at marine hydrothermal vents. mSystems 5:e00553–19

21. Barco RA, Merino N, Lam B, Budnik B, Kaplan M, Wu F, Amend JP, Nealson KH, Emerson D. 2024. Comparative proteomics of a versatile, marine, iron-oxidizing chemolithoautotroph. Environ Microbiol 26:e16632.

22. Zhou N, Kupper RJ, Catalano JG, Thompson A, Chan CS. 2022. Biological oxidation of Fe(II)-bearing smectite by microaerophilic iron oxidizer *Sideroxydans lithotrophicus* using dual Mto and Cyc2 iron oxidation pathways. Environ Sci Technol 56:17443–17453.

23. Keffer JL, Zhou N, Rushworth DD, Yu Y, Chan CS. 2024. Microbial magnetite oxidation via MtoAB porin-multiheme cytochrome complex in *Sideroxydans lithotrophicus* ES-1. bioRxiv 10.1101/2024.09.20.614158. In press at AEM.

24. He S, Barco RA, Emerson D, Roden EE. 2017. Comparative genomic analysis of neutrophilic iron(II) oxidizer genomes for candidate genes in extracellular electron transfer. Front Microbiol 8:1584.

25. Zhou N. 2022. The molecular mechanisms and organic effects on microaerophilic Fe(II) oxidation by *Sideroxydans Lithotrophicus* ES-1. Doctoral dissertation, University of Delaware.

26. Cooper RE, Wegner C-E, Kügler S, Poulin RX, Ueberschaar N, Wurlitzer JD, Stettin D, Wichard T, Pohnert G, Küsel K. 2020. Iron is not everything: unexpected complex metabolic responses between iron-cycling microorganisms. The ISME Journal 14:2675– 2690.

27. Ohta Y, Nogi Y, Miyazaki M, Li Z, Hatada Y, Ito S, Horikoshi K. 2004. Enzymatic properties and nucleotide and amino acid sequences of a thermostable β-agarase from the novel marine isolate, JAMB-A94. Biosci Biotechnol Biochem 68:1073–1081.

28. Bannikova GE, Lopatin SA, Varlamov VP, Kuznetsov BB, Kozina IV, Miroshnichenko ML, Chernykh NA, Turova TP, Bonch-Osmolovskaya EA. 2008. The thermophilic bacteria hydrolyzing agar: Characterization of thermostable agarase. Appl Biochem Microbiol 44:366–371.

29. Badger MR, Bek EJ. 2008. Multiple Rubisco forms in proteobacteria: their functional significance in relation to CO_2_ acquisition by the CBB cycle. J Exp Bot 59:1525–1541.

30. Beckwith CR, Edwards MJ, Lawes M, Shi L, Butt JN, Richardson DJ, Clarke TA. 2015. Characterization of MtoD from *Sideroxydans lithotrophicus*: a cytochrome c electron shuttle used in lithoautotrophic growth. Front Microbiol 6:332.

31. Jain A, Kalb MJ, Gralnick JA. 2022. Reconstructing electron transfer components from an Fe(II) oxidizing bacterium. Microbiology 168:001240.

32. Jain A, Coelho A, Madjarov J, Paquete CM, Gralnick JA. 2022. Evidence for quinol oxidation activity of ImoA, a novel NapC/NirT family protein from the neutrophilic Fe(II)-oxidizing bacterium *Sideroxydans lithotrophicus* ES-1. mBio 13:e02150–22.

33. Liu J, Wang Z, Belchik SM, Edwards MJ, Liu C, Kennedy DW, Merkley ED, Lipton MS, Butt JN, Richardson DJ, Zachara JM, Fredrickson JK, Rosso KM, Shi L. 2012. Identification and Characterization of MtoA: a decaheme c-type cytochrome of the neutrophilic Fe(II)-oxidizing bacterium *Sideroxydans lithotrophicus* ES-1. Front Microbio 3:37.

34. Keffer JL, McAllister SM, Garber AI, Hallahan BJ, Sutherland MC, Rozovsky S, Chan CS. 2021. Iron oxidation by a fused cytochrome-porin common to diverse iron-oxidizing bacteria. mBio 12:e01074–21.

35. Kappler A, Bryce C, Mansor M, Lueder U, Byrne JM, Swanner ED. 2021. An evolving view on biogeochemical cycling of iron. Nat Rev Microbiol 19:360–374.

36. Huang J, Jones A, Waite TD, Chen Y, Huang X, Rosso KM, Kappler A, Mansor M, Tratnyek PG, Zhang H. 2021. Fe(II) Redox chemistry in the environment. Chem Rev 121: 8161–8233.

37. Levar CE, Rollefson DR, Bond. 2012. Microbial Metal Respiration: From Geochemistry to Potential Applications. Springer Berlin Heidelberg p 29–48.

38. Joshi K, Chan CH, Bond DR. 2021. Geobacter sulfurreducens inner membrane cytochrome CbcBA controls electron transfer and growth yield near the energetic limit of respiration. Mol Microbiol 116:1124–1139.

39. Rohwer RR, Kirkpatrick M, Garcia SL, Kellom M, McMahon KD, Baker BJ. 2025. Two decades of bacterial ecology and evolution in a freshwater lake. Nat Microbiol 10:246–257.

40. Oliver T, Varghese N, Roux S, Schulz F, Huntemann M, Clum A, Foster B, Foster B, Riley R, LaButti K, Egan R, Hajek P, Mukherjee S, Ovchinnikova G, Reddy TBK, Calhoun S, Hayes RD, Rohwer RR, Zhou Z, Daum C, Copeland A, Chen I-MA, Ivanova NN, Kyrpides NC, Mouncey NJ, del Rio TG, Grigoriev IV, Hofmeyr S, Oliker L, Yelick K, Anantharaman K, McMahon KD, Woyke T, Eloe-Fadrosh EA. 2024. Coassembly and binning of a twenty-year metagenomic time-series from Lake Mendota. Sci Data 11:966.

41. Buck M, Garcia SL, Fernandez L, Martin G, Martinez-Rodriguez GA, Saarenheimo J, Zopfi J, Bertilsson S, Peura S. 2021. Comprehensive dataset of shotgun metagenomes from oxygen stratified freshwater lakes and ponds. Sci Data 8:131.

42. Hädrich A, Heuer VB, Herrmann M, Hinrichs K-U, Küsel K. 2012. Origin and fate of acetate in an acidic fen. FEMS Microbiol Ecol 81:339–354.

43. Roden EE, Wetzel RG. 1996. Organic carbon oxidation and suppression of methane production by microbial Fe(III) oxide reduction in vegetated and unvegetated freshwater wetland sediments. Limnol Oceanogr 41:1733–1748.

44. Roden EE, Wetzel RG. 2003. Competition between Fe(III)-reducing and methanogenic bacteria for acetate in iron-rich freshwater sediments. Microb Ecol 45:252–258.

45. Reiche M, Torburg G, Küsel K. 2008. Competition of Fe(III) reduction and methanogenesis in an acidic fen. FEMS Microbiol Ecol 65:88–101.

46. Teh YA, Dubinsky EA, Silver WL, Carlson CM. 2008. Suppression of methanogenesis by dissimilatory Fe(III)-reducing bacteria in tropical rain forest soils: implications for ecosystem methane flux. Glob Change Biol 14:413–422.

47. Stookey LL. 1970. Ferrozine—a new spectrophotometric reagent for iron. Anal Chem 42:779–781.

48. Riemer J, Hoepken HH, Czerwinska H, Robinson SR, Dringen R. 2004. Colorimetric ferrozine-based assay for the quantitation of iron in cultured cells. Anal Biochem 331:370– 375.

49. Babraham Bioinformatics. FastQC a quality control tool for high throughput sequence data. http://www.bioinformatics.babraham.ac.uk/projects/fastqc/

50. Krueger F. Trim Galore. The Babraham Institute. https://github.com/FelixKrueger/TrimGalore

51. Martin M. 2011. Cutadapt removes adapter sequences from high-throughput sequencing reads. 1. EMBnet.journal 17:10–12.

52. Kopylova E, Noé L, Touzet H. 2012. SortMeRNA: fast and accurate filtering of ribosomal RNAs in metatranscriptomic data. Bioinformatics 28:3211–3217.

53. Langmead B, Salzberg SL. 2012. Fast gapped-read alignment with Bowtie 2. Nat Methods 9:357–359.

54. Li H, Handsaker B, Wysoker A, Fennell T, Ruan J, Homer N, Marth G, Abecasis G, Durbin R, 1000 Genome Project Data Processing Subgroup. 2009. The sequence alignment/map format and SAMtools. Bioinforma Oxf Engl 25:2078–2079.

55. Anders S, Pyl PT, Huber W. 2015. HTSeq—a Python framework to work with high-throughput sequencing data. Bioinformatics 31:166–169.

56. R Core Team. 2024. R: A Language and Environment for Statistical Computing. R Foundation for Statistical Computing, Vienna, Austria.

57. Posit team. 2023. RStudio: Integrated Development Environment for R (1.4.1717). Posit Software, PBC, Boston, MA.

58. Love MI, Huber W, Anders S. 2014. Moderated estimation of fold change and dispersion for RNA-seq data with DESeq2. Genome Biol 15:550.

59. Gilchrist CLM, Booth TJ, van Wersch B, van Grieken L, Medema MH, Chooi Y-H. 2021. cblaster: a remote search tool for rapid identification and visualization of homologous gene clusters. Bioinforma Adv 1:vbab016.

60. Gilchrist CLM, Chooi Y-H. 2021. clinker & clustermap.js: automatic generation of gene cluster comparison figures. Bioinformatics 37:2473–2475.

61. Yu NY, Wagner JR, Laird MR, Melli G, Rey S, Lo R, Dao P, Sahinalp SC, Ester M, Foster LJ, Brinkman FSL. 2010. PSORTb 3.0: improved protein subcellular localization prediction with refined localization subcategories and predictive capabilities for all prokaryotes. Bioinformatics 26:1608–1615.

62. Shen H-B, Chou K-C. 2010. Gneg-mPLoc: A top-down strategy to enhance the quality of predicting subcellular localization of Gram-negative bacterial proteins. J Theor Biol 264:326–333.

63. Bernhofer M, Dallago C, Karl T, Satagopam V, Heinzinger M, Littmann M, Olenyi T, Qiu J, Schütze K, Yachdav G, Ashkenazy H, Ben-Tal N, Bromberg Y, Goldberg T, Kajan L, O’Donoghue S, Sander C, Schafferhans A, Schlessinger A, Vriend G, Mirdita M, Gawron P, Gu W, Jarosz Y, Trefois C, Steinegger M, Schneider R, Rost B. 2021. PredictProtein - predicting protein structure and function for 29 years. Nucleic Acids Res 49:W535–W540.

64. Jones P, Binns D, Chang H-Y, Fraser M, Li W, McAnulla C, McWilliam H, Maslen J, Mitchell A, Nuka G, Pesseat S, Quinn AF, Sangrador-Vegas A, Scheremetjew M, Yong S-Y, Lopez R, Hunter S. 2014. InterProScan 5: genome-scale protein function classification. Bioinformatics 30:1236–1240.

65. Blum M, Chang H-Y, Chuguransky S, Grego T, Kandasaamy S, Mitchell A, Nuka G, Paysan-Lafosse T, Qureshi M, Raj S, Richardson L, Salazar GA, Williams L, Bork P, Bridge A, Gough J, Haft DH, Letunic I, Marchler-Bauer A, Mi H, Natale DA, Necci M, Orengo CA, Pandurangan AP, Rivoire C, Sigrist CJA, Sillitoe I, Thanki N, Thomas PD, Tosatto SCE, Wu CH, Bateman A, Finn RD. 2021. The InterPro protein families and domains database: 20 years on. Nucleic Acids Res 49:D344–D354.

