## Supplemental Figures and Table S3 for "An organotrophic *Sideroxydans* reveals potential iron oxidation marker genes"

Number of pages: 6
Number of figures: 2
Number of tables: 1 (Table S3)

Tables S1 and S2 are available as xls files.

A)

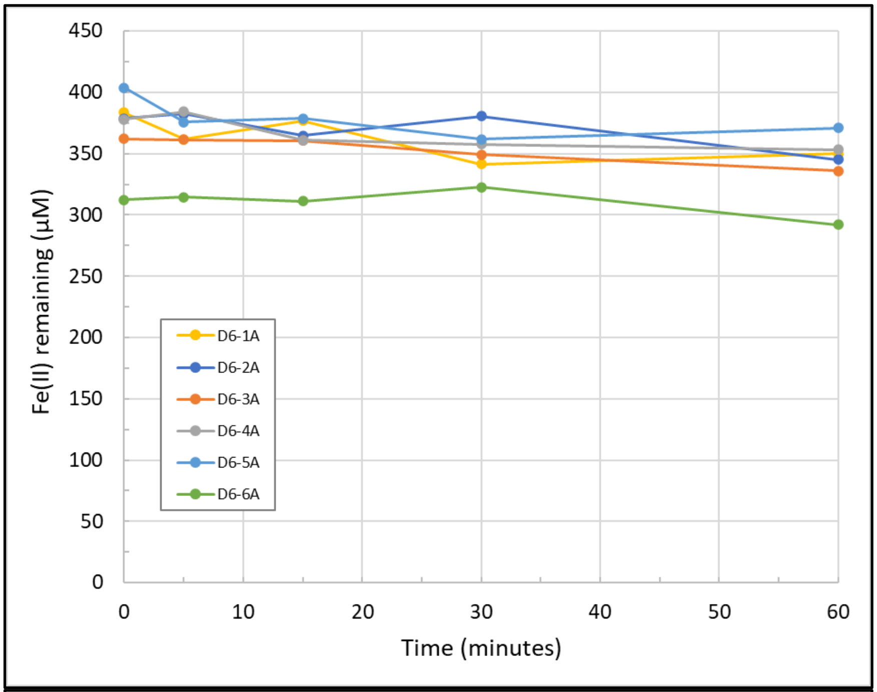
450
400
350
300
250
200
150
100
50
0
0
10
20
30
40
50
60
Time (minutes)
Fe(II) remaining (µM)
D6-1A
D6-2A
D6-3A
D6-4A
D6-5A
D6-6A

B)

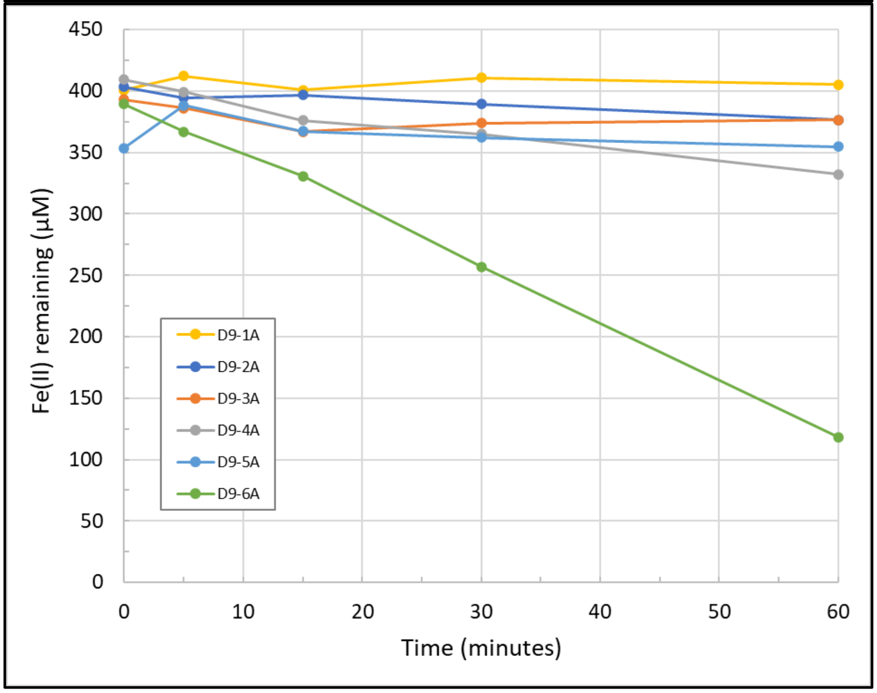
450
400
350
300
250
200
150
100
50
0
0
10
20
30
40
50
60
Time (minutes)
Fe(II) remaining (µM)
D9-1A
D9-2A
D9-3A
D9-4A
D9-5A
D9-6A

C)

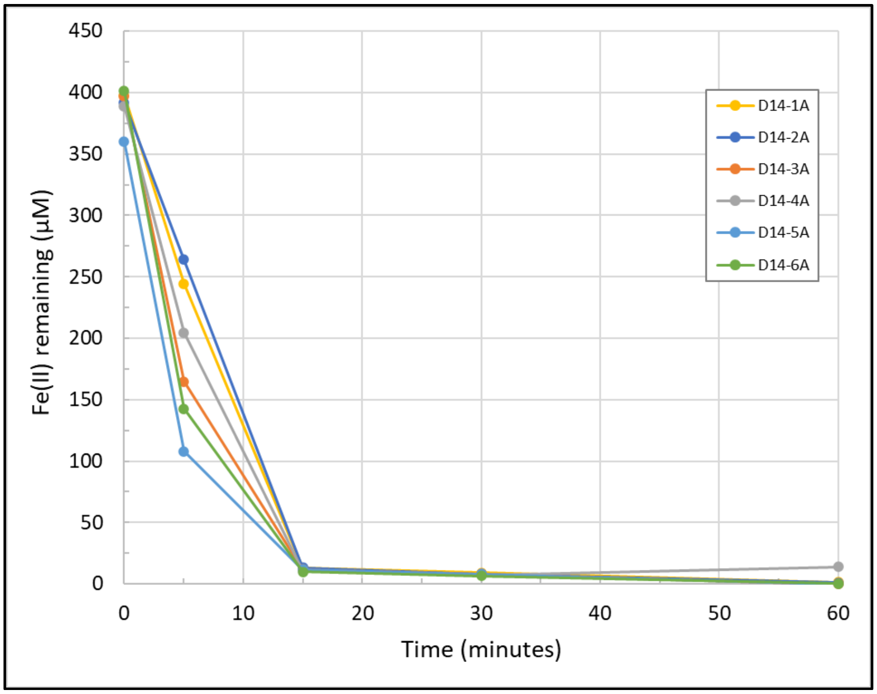


**FIG S1.** Iron oxidation response of individual lactate-grown bottles during the Fe(II)-spike experiments on A) Day 6, B) Day 9, and C) Day 14.

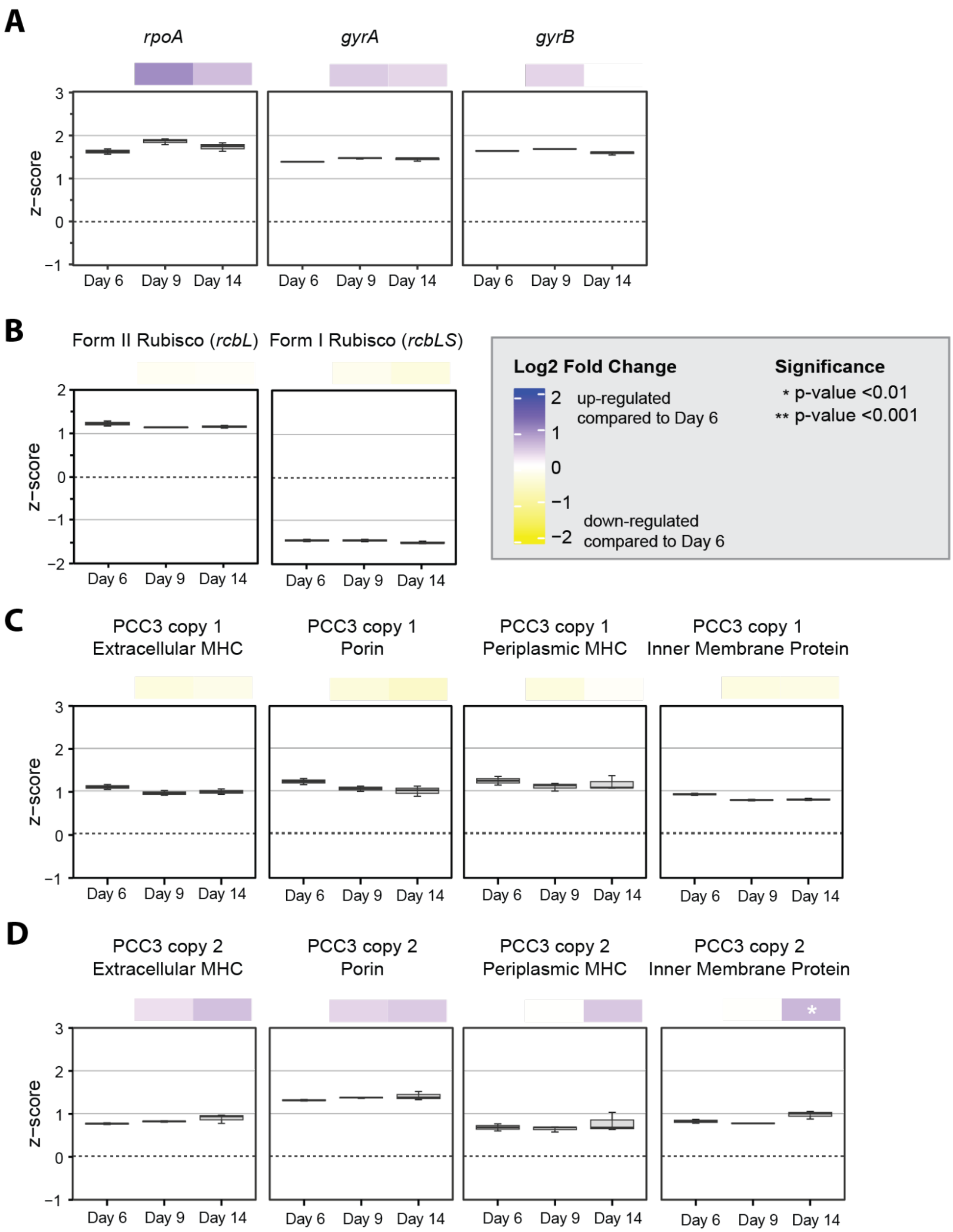


**FIG S2.** Boxplots of z-score normalized gene expression for each time point with heatmaps showing the Log2 Fold Change in expression between Day 9 and Day 14 versus Day 6. Expression is represented by Z-score, which shows the expression of each gene relative to the mean gene expression (0) at the time point. Z-score was calculated from rlog normalized count data from DESeqs2. Data is shown for A) housekeeping genes, B) Form I and Form II RuBisCo, C) PCC3 copy 1, and PCC3 copy 2. Error bars represent the spread of the interquartile range.

**TABLE S3.** Existing evidence for the involvement of various genes and proteins in iron oxidation and/or extracellular electron transport.

| Gene(s)/ complex | Evidence for EET and/or Fe oxidation | Reference |
|---|---|---|
| ***grcJ/* GrcJ** | ● $GrcJ_{TAG\text{-}1}$ upregulated during iron oxidation<br>● *grcJ*$_{TAG\text{-}1}$ expression increased over time when cultured on Fe | Barco et al. 2024 (1)<br>Barco et al. 2024 (1) |
| ***irc*** | ● *ircABCD*$_{ES\text{-}1}$ upregulated under Fe-oxidizing v. $Na_2S_2O_3$-oxidizing conditions (referred to as Slit_1321-Slit_1324) | Zhou et al. 2021 (2) |
| **PCC3** | ● Identifies PCC3 as a set of genes encoding porin-cytochrome complexes similar to *mtoAB/pioAB* and are present in many FeOB<br>● Heterologous expression of inner membrane and periplasmic components of $PCC3_{ES\text{-}1}$ shows it is likely modular and able to transport electrons | He et al. 2017 (3)<br>Jain, et al. 2022 (4) |
| ***mtoD*** | ● Biochemical, biophysical, and crystallographic characterization of MtoD indicates it can shuttle electrons through the periplasm<br>● Heterologous expression of *mtoD*$_{ES\text{-}1}$ shows it is likely modular and involved in EET | Beckwith et al. 2015 (5)<br>Jain, et al. 2022 (4) |
| ***cymA/ imoA*** | ● Homology and predicted localization of $CymA_{ES\text{-}1}$ make it a potential pathway component for EET<br>● CymA/ImoA may interact with the quinone pool<br>● Heterologous expression of *imoA*$_{ES\text{-}1}$ shows involvement in EET<br>● CymA/$ImoA_{ES\text{-}1}$ only detected when cells were cultured on solid Fe (magnetite), not dissolved Fe(II) | Liu et al. 2012 (6)<br>Jain et al. 2022 (7)<br>Jain, et al. 2022 (4)<br>Keffer et al. 2024 (8) |
| ***cyc2*** | ● $Cyc2_{PV\text{-}1}$ can oxidize Fe(II)<br>● $Cyc2_{TAG\text{-}1}$ highly expressed under $H_2$- and Fe(II)-oxidizing conditions<br>● *cyc2*$_{ES\text{-}1}$ highly expressed overall with different expression patterns for various gene copies<br>● *cyc2*$_{ES\text{-}1}$ highly expressed on dissolved Fe(II) and solid Fe(II)-bearing smectite clay<br>● $Cyc2_{ES\text{-}1}$ highly expressed on both dissolved and solid Fe substrates<br>● *cyc2* is highly expressed in many Zetaproteobacteria and expression becomes even higher after an addition of dissolved Fe(II)<br>● *cyc2*$_{CL21}$ highly expressed on Fe(0) (supplemental data) | Keffer et al. 2021 (9)<br>Barco et al. 2024 (1)<br>Zhou et al. 2021 (2)<br>Zhou et al. 2022 (10)<br>Keffer et al. 2024 (8)<br>McAllister, et al. 2020 (11)<br>McAllister, et al. 2020 (12)<br>Cooper, et al. 2020 (13) |
| ***mtoAB*** | ● MtoA can oxidize Fe(II). MtoA also rescues some iron reduction function in *Shewanella oneidensis* MR-1<br>● *mtoA*$_{CL21}$ highly expressed on Fe(0) in mono-culture and in co-culture with *S. oneidensis* MR-1<br>● $MtoA_{ES\text{-}1}$ is not highly expressed on dissolved Fe(II)<br>● *mtoA*$_{ES\text{-}1}$ expression is upregulated on solid Fe-bearing smectite v. dissolved Fe(II)<br>● $MtoA_{ES\text{-}1}$ was only detected when cells were cultured on solid Fe (magnetite), not dissolved Fe(II) | Liu et al. 2012 (6)<br>Cooper, et al. 2020 (13)<br>Zhou et al. 2021 (2)<br>Zhou et al. 2022 (10)<br>Keffer et al. 2024 (8) |

### References


1. Barco RA, Merino N, Lam B, Budnik B, Kaplan M, Wu F, Amend JP, Nealson KH, Emerson D. 2024. Comparative proteomics of a versatile, marine, iron-oxidizing chemolithoautotroph. Environmental Microbiology 26:e16632.
2. Zhou N, Keffer JL, Polson SW, Chan CS. 2021. Unraveling Fe(II)-Oxidizing Mechanisms in a Facultative Fe(II) Oxidizer, *Sideroxydans lithotrophicus* Strain ES-1, via Culturing, Transcriptomics, and Reverse Transcription-Quantitative PCR. Applied and Environmental Microbiology 88:e01595-21.
3. He S, Barco RA, Emerson D, Roden EE. 2017. Comparative Genomic Analysis of Neutrophilic Iron(II) Oxidizer Genomes for Candidate Genes in Extracellular Electron Transfer. Front Microbiol 8:1584.
4. Jain A, Kalb MJ, Gralnick JA. 2022. Reconstructing electron transfer components from an Fe(II) oxidizing bacterium. Microbiology 168:001240.
5. Beckwith CR, Edwards MJ, Lawes M, Shi L, Butt JN, Richardson DJ, Clarke TA. 2015. Characterization of MtoD from *Sideroxydans lithotrophicus*: a cytochrome c electron shuttle used in lithoautotrophic growth. Front Microbiol 6:332.
6. Liu J, Wang Z, Belchik SM, Edwards MJ, Liu C, Kennedy DW, Merkley ED, Lipton MS, Butt JN, Richardson DJ, Zachara JM, Fredrickson JK, Rosso KM, Shi L. 2012. Identification and Characterization of MtoA: A Decaheme c-Type Cytochrome of the Neutrophilic Fe(II)-Oxidizing Bacterium *Sideroxydans lithotrophicus* ES-1. Front Microbio 3:37.
7. Jain A, Coelho A, Madjarov J, Paquete CM, Gralnick JA. 2022. Evidence for Quinol Oxidation Activity of ImoA, a Novel NapC/NirT Family Protein from the Neutrophilic Fe(II)-Oxidizing Bacterium *Sideroxydans lithotrophicus* ES-1. mBio 13:e02150-22.
8. Keffer JL, Zhou N, Rushworth DD, Yu Y, Chan CS. 2024. Microbial magnetite oxidation via MtoAB porin-multiheme cytochrome complex in Sideroxydans lithotrophicus ES-1. bioRxiv https://doi.org/10.1101/2024.09.20.614158. In press at Appl. Env. Microbiol.
9. Keffer JL, McAllister SM, Garber AI, Hallahan BJ, Sutherland MC, Rozovsky S, Chan CS. 2021. Iron Oxidation by a Fused Cytochrome-Porin Common to Diverse Iron-Oxidizing Bacteria. mBio 12:e01074-21.
10. Zhou N, Kupper RJ, Catalano JG, Thompson A, Chan CS. 2022. Biological Oxidation of Fe(II)-Bearing Smectite by Microaerophilic Iron Oxidizer *Sideroxydans lithotrophicus* Using Dual Mto and Cyc2 Iron Oxidation Pathways. Environ Sci Technol https://doi.org/10.1021/acs.est.2c05142.
11. McAllister SM, Polson SW, Butterfield DA, Glazer BT, Sylvan JB, Chan CS. 2020. Validating the Cyc2 Neutrophilic Iron Oxidation Pathway Using Meta-omics of Zetaproteobacteria Iron Mats at Marine Hydrothermal Vents. mSystems 5:e00553-19, /msystems/5/1/msys.00553-19.atom.
12. McAllister SM, Vandzura R, Keffer JL, Polson SW, Chan CS. 2020. Aerobic and anaerobic iron oxidizers together drive denitrification and carbon cycling at marine iron-rich hydrothermal vents. The ISME Journal 5:1271-1286.
13. Cooper RE, Wegner C-E, Kügler S, Poulin RX, Ueberschaar N, Wurlitzer JD, Stettin D, Wichard T, Pohnert G, Küsel K. 2020. Iron is not everything: unexpected complex metabolic responses between iron-cycling microorganisms. The ISME Journal 14:2675-2690.
